# Neural correlates of the shamanic state of consciousness

**DOI:** 10.1101/2020.07.20.212522

**Authors:** Emma R. Huels, Hyoungkyu Kim, UnCheol Lee, Tarik Bel-Bahar, Angelo Colmenero, Amanda Nelson, Stefanie Blain-Moraes, George A. Mashour, Richard E. Harris

## Abstract

Despite the use of shamanism as a healing practice for several millennia, few empirical studies of the shamanic state of consciousness exist. We investigated the neural correlates of shamanic trance using high-density electroencephalography (EEG) in 24 shamanic practitioners and 24 healthy controls during rest, shamanic drumming, and classical music listening, followed by a validated assessment of altered states of consciousness. EEG data were used to assess changes in absolute power, connectivity, signal diversity, and criticality, which were correlated with assessment measures. We also compared assessment scores to those of individuals in a previous study under the influence of psychedelics. Shamanic practitioners were significantly different from controls in several domains of altered states of consciousness, with scores comparable to or exceeding that of healthy volunteers under the influence of psychedelics. Practitioners also displayed increased gamma power during drumming that positively correlated with elementary visual alterations. Furthermore, shamanic practitioners had decreased low alpha and increased low beta connectivity during drumming and classical music, and decreased neural signal diversity in the gamma band during drumming that inversely correlated with insightfulness. Finally, criticality in practitioners was increased during drumming in the low and high beta and gamma bands, with increases in the low beta band correlating with complex imagery and elementary visual alterations. These findings suggest that psychedelic drug-induced and non-pharmacologic alterations in consciousness have overlapping phenomenal traits but are distinct states of consciousness, as reflected by the unique brain-related changes during shamanic trance compared to previous literature investigating the psychedelic state.

## Introduction

Shamanism, an ancient spiritual practice of indigenous cultures throughout the world, has been used for physical, psychological, and spiritual healing since the Paleolithic era(1). Although wide variations of shamanic practice exist across history and cultures(2), core shamanic features are present in modern-day religions and Western neo-shamanism(3). Despite its universality in ancient civilizations and increasing popularity in the present day, empirical studies of shamanic practice remain scarce.

During this form of spiritual practice, the shaman (in cases of indigenous tribes) or shamanic practitioner (in cases of neo-shamanism in the western world) enters an altered state of consciousness, termed the shamanic state of consciousness(4), through pharmacologic or non-pharmacologic means(5). Pharmacologic means of entering this state typically involve ingesting psychedelic substances, such as Ayahuasca, whereas non-pharmacologic means usually involve listening to a slow (2 - 4 Hz), repetitive rhythm, such as a drum or rattle, and dancing(6). During the shamanic state of consciousness, shamanic practitioners enter a trance state, which has significant anecdotal overlap with the experience of individuals under the influence of psychedelic substances. Specifically, both groups may encounter mystical experiences or entities, feelings of disembodiment or flight, or feelings of ego dissolution(7).

Despite widespread anthropological research of shamanism, only two neuroimaging studies have investigated brain activity during shamanic trance. Using fMRI, Hove et al. found increased hub activity in key regions in the default mode and salience networks during the shamanic state(8). Functional connectivity between these regions was also increased during trance, suggesting a “merging” of internal and external sensory processing. Research using electroencephalographic activity during the shamanic state of consciousness revealed increased low (13-20 Hz) and high (21-50 Hz) beta activity, with low beta localized to frontal regions and high beta localized to parietal regions(9). Although these studies have contributed to our knowledge of the shamanic state, the Hove et al. trial lacked non-shamanic practitioner controls and Flor-Henry et al. examined data from only a single shamanic practitioner. Further, these past studies did not correlate neurobiological metrics with the subjective experiences of this state.

To address these gaps, we investigated the shamanic state of consciousness using a validated assessment of altered states of consciousness and high-density electroencephalography (EEG) in both experienced shamanic practitioners and naïve control participants. While spectral analyses remain standard in many EEG studies, recent investigations regarding altered states of consciousness have utilized other computational measures to characterize brain activity, such as functional connectivity(10, 11), signal diversity(12–14), and criticality(15–18). Each of these measures informs on various attributes of the neural signal—functional relationships, complexity (i.e., number of different patterns in the signal), and the balance between order and disorder, respectively—that better elucidate the neural correlates underlying brain states. Thus, we characterized EEG changes during shamanic drumming using spectral analysis, connectivity, signal diversity, and criticality, and determined their relationship to changes in a variety of domains used to characterize altered states of consciousness. Further, we compared these self-reported experiences during shamanic trance to the altered state of consciousness induced by psychedelic substances.

## Results

### Shamanic practitioners enter an altered state of consciousness during drumming

We first evaluated differences in OAV domain scores to verify that shamanic practitioners entered an altered state of consciousness. Shamanic practitioners had significantly greater scores than control participants in 8 of the 11 domains during shamanic healing, including complex imagery (Z = 4.22, *p* < .001, 95% CI [0.30, 0.57]), experience of unity (Z = 4.04, *p* < .001, 95% CI [0.33, 0.65]), spiritual experience (Z = 4.87, *p* < .001, 95% CI [0.33, 0.60]), blissful state (Z = 4.39, *p* < .001, 95% CI [0.30, 0.60]), disembodiment (Z = 4.4, *p* < .001, 95% CI [0.23, 0.67]), insightfulness (Z = 4.06, *p* < .001, 95% CI [0.23, 0.57]), elementary visual alterations (Z = 2.43, *p* = .0159, 95% CI [0.033, 0.40]), and changed meaning of percepts (Z = 3.22, *p* = .00136, 95% CI [0.033, 0.40)(**Figure 1A**). These differences did not exist when comparing OAV domain scores during the classical music experiment (**Figure 1B**). Furthermore, shamanic practitioners were significantly different in all of the above domains except elementary visual alterations when comparing their scores during shamanic drumming to classical music (Z = 2.98 to Z = 3.62, *p* < .01 to *p* < .001), indicating a majority of the increased scores were specific to the shamanic drumming experiment. Finally, shamanic practitioners had increased scores in audio-visual synesthesia (Z = -2.35, *p* = .0199, 95% CI [-0.42, -0.033]) and control participants had increased scores in blissful state (Z = -2.42, p = .017, 95% CI [-0.23, -0.017]) during classical music listening compared to drumming.

**Figure 1.**
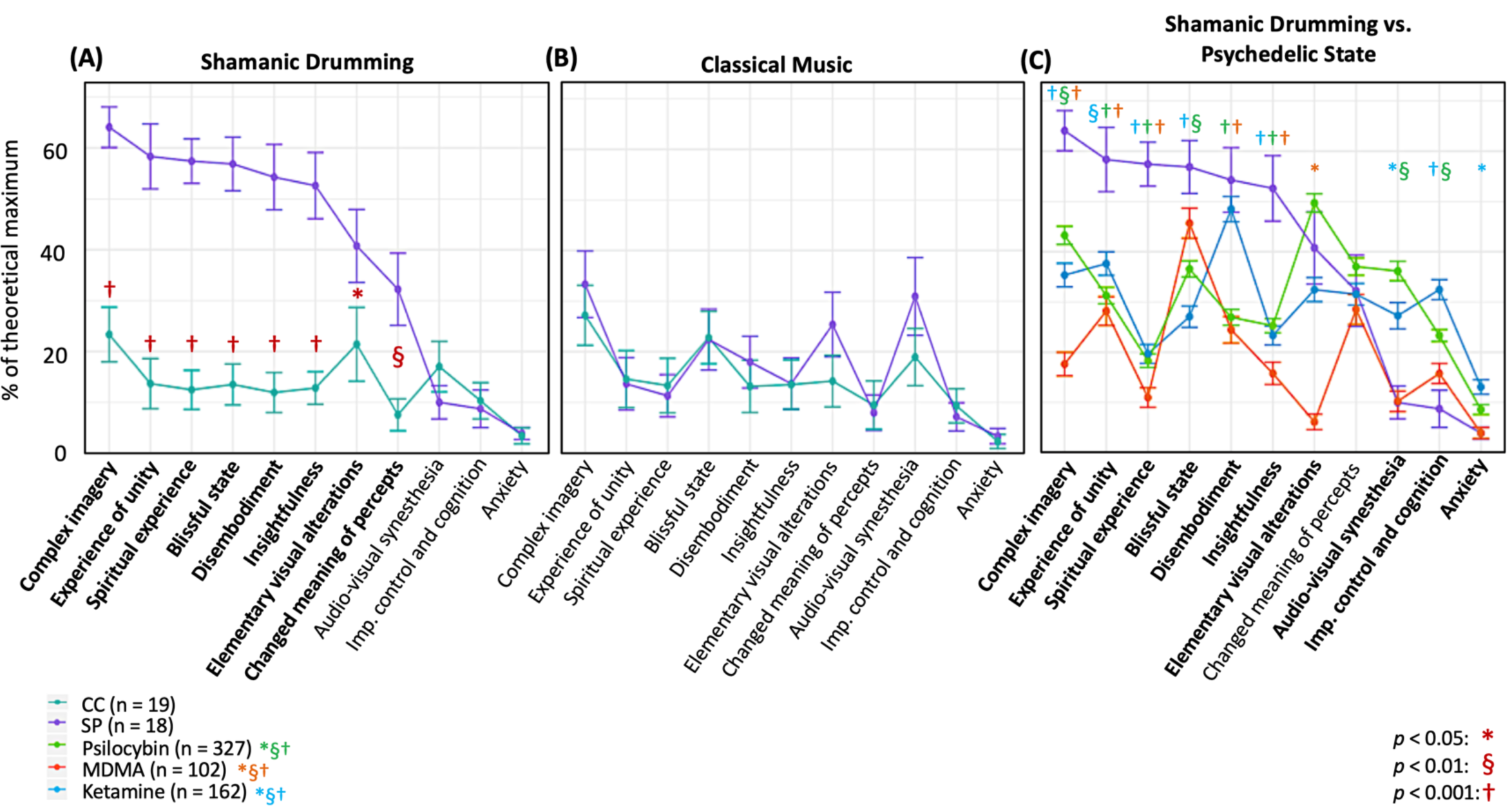
Shamanic practitioners enter an altered state of consciousness during shamanic drumming. **(A)** Shamanic Practitioners (SP, purple, n = 18) were significantly different from control participants (CC, green, n = 19) during drumming in 8 of the 11 OAV domains. Each point represents the mean of all SP or CC for the percent of the theoretical scale maximum for that domain. Error bars represent the standard error of the mean. **(B)** SP did not differ from CC during classical music listening on any of the domains. **(C)** We replotted the OAV mean and standard error for healthy controls taking psilocybin (green, n = 327), ketamine (blue, n = 162), or MDMA (orange, n = 102) from Studerus, Gamma, & Vollenweider (2010) and compared these scores to those acquired from SP. Overall, SP had increased scores in several domains compared to all three drug conditions, as well as similar scores in the remaining domains compared to various drug conditions.

### Shamanic practitioners experience an altered state of consciousness similar to yet distinct from the psychedelic state

Given the overlap between the shamanic state of consciousness and the psychedelic experience, we thought it would be informative to compare the OAV domain scores from shamanic practitioners with those of individuals under the influence of psychedelic substances. Studerus et al. pooled OAV scores from healthy individuals under the influence of psilocybin, ketamine, and MDMA(19). With author permission, we replotted the mean and standard error of the mean (SEM) for each drug condition from Studerus et al. with our data from shamanic practitioners, and compared each of the domains for each drug condition to shamanic practitioners using unpaired t-tests. As can be seen in **Figure 1C**, shamanic practitioners experienced several domains at or above the level of individuals under the influence of various psychedelics. Specifically, the domains of complex imagery (ketamine: *p* < .001; psilocybin: *p* = .0077; MDMA: *p* < .001), experience of unity (ketamine: *p* = .0052; psilocybin: *p* < .001; MDMA: *p* < .001), spiritual experience (ketamine: *p* < .001; psilocybin: *p* < .001; MDMA: *p* < .001), and insightfulness (ketamine: *p* < .001; psilocybin: *p* < .001; MDMA: *p* < .001) appear to be greater for shamanic practitioners during journeying compared to healthy controls under any of these very potent psychedelic compounds. However, shamanic practitioners were similar to MDMA in the domains of blissful state (ketamine: *p* < .001; psilocybin: *p* = .0036; MDMA: *p* = .14), audio-visual synesthesia (ketamine: *p* = .031; psilocybin: *p* = .0017; MDMA: *p* = .96), and impaired control and cognition (ketamine: *p* < .001; psilocybin: *p* = .0045; MDMA: *p* = .16), but comparable to ketamine in the domain of disembodiment (ketamine: *p* = .46; psilocybin: *p* < .001; MDMA: *p* < .001), and ketamine and psilocybin in elementary visual alterations (ketamine: *p* =. 28; psilocybin: *p* = .27; MDMA: *p* < .001). Additionally, individuals under the influence of ketamine had significantly higher anxiety scores (ketamine: *p* = .037; psilocybin: *p* = .27; MDMA: *p* = .99) compared to shamanic practitioners during drumming. Finally, practitioners were similar to all three drugs in the domain of changed meaning of percepts (ketamine: *p* = .92; psilocybin: *p* = .52; MDMA: *p* = .63). Additional statistical output measures are detailed in **Supplemental Table 1**.

### Absolute gamma power increases during drumming in shamanic practitioners and correlates with elementary visual alterations

Next, we assessed differences in absolute power between shamanic practitioners and controls during eyes closed, classical music, and drumming (**Figure 2A**). There was a statistically significant interaction between group and condition on gamma power (Group x Condition: *F*(1.7, 59.1) = 6.3, *p* = .005). Post-hoc exploratory t-tests revealed shamanic practitioners had significantly greater gamma power during drumming compared to controls (*t*(29) = 2.16, *p* = .039, 95% CI [0.040, 1.43])(**Figure 2B**). Further, gamma power during drumming positively correlated with the degree of elementary visual alterations (*r*_*s*_ = 0.52, *p* = .025)(**Figure 2C**) in shamanic practitioners.

**Figure 2.**
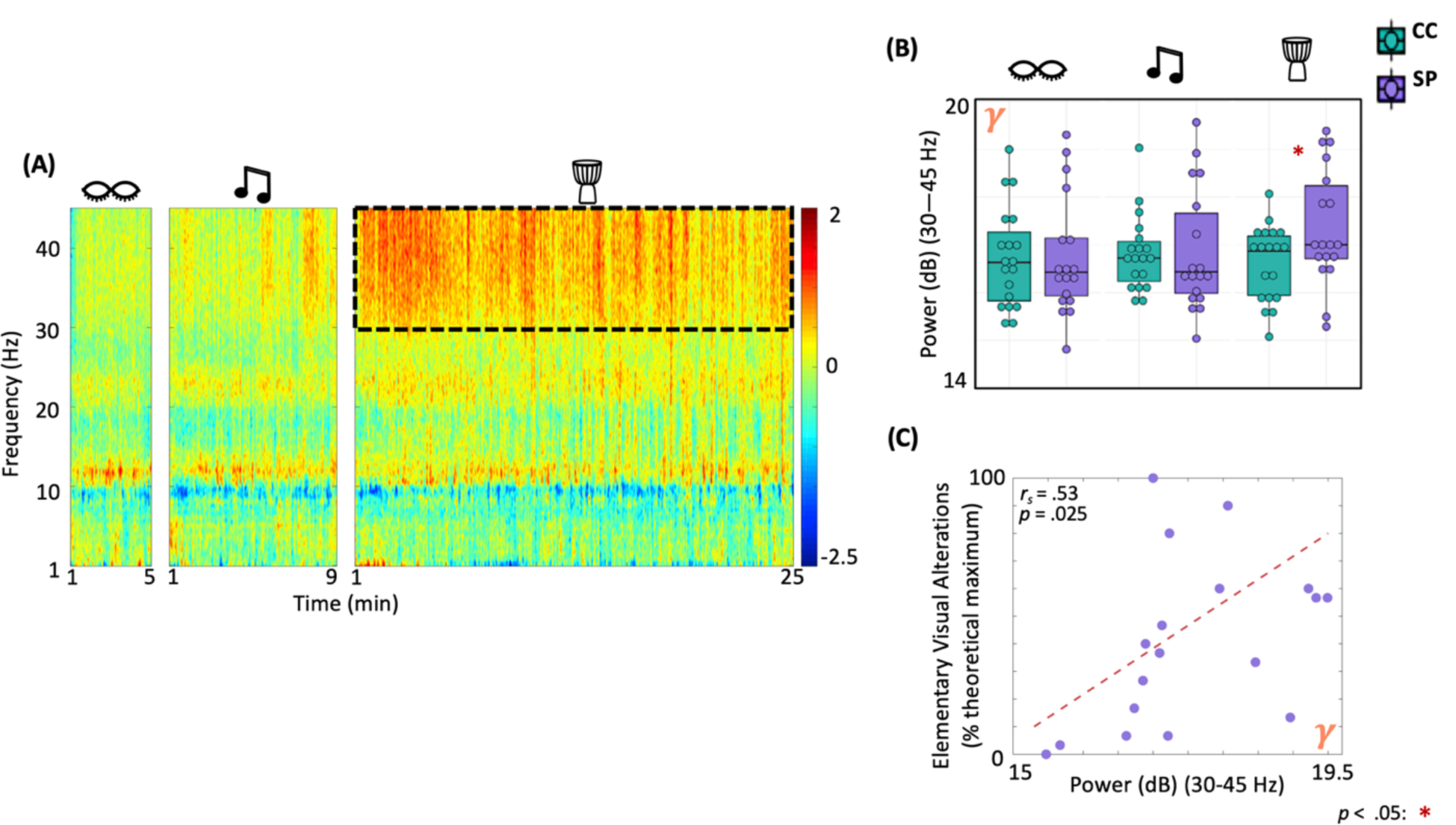
Gamma power is greater in shamanic practitioners during drumming and correlates with elementary visual alterations. **(A)** Power spectrogram comparing the difference in power across frequencies (1-45 Hz) between shamanic practitioners (SP; n = 18) and control participants (CC; n = 19) during eyes closed, classical music, and drumming periods. Warmer colors represent greater power in SP and cooler colors indicate more power in CC. The dotted line represents the gamma frequency band (30-45 Hz), which was significantly different between groups during shamanic drumming. **(B)** Box plots representing the mean absolute power of SP (purple) or CC (green) during eyes closed, drumming, or music. Whiskers represent the lower (25%) and upper (75%) quartiles and the center band of each box represents the median for the group. SP had significantly greater gamma power during the drumming period compared to CC. Significance is indicated by the key in the lower righthand corner of the figure. **(C)** We correlated absolute gamma power with each of the eight ASC domain scores that were significantly different between groups during drumming in **Figure 1A**, revealing a positive correlation between absolute gamma power and the degree of elementary visual alterations.

### Shamanic practitioners have altered functional connectivity in the low alpha and beta bands during drumming

We also examined differences in functional connectivity within each state in shamanic practitioners and control participants (**Figure 3A**). There was a significant main effect of group on low alpha (*F*(1,35) = 5.6, *p* = .024) and low beta (*F*(1,35) = 4.41, *p* = .043) connectivity. Post-hoc t-tests revealed that shamanic practitioners had significantly decreased low alpha connectivity during drumming (*t*(27.2) = -2.45, *p* = .021, 95% CI [0.00731, 0.0828]) and music (*t*(27.8) = -2.22, *p* = .035, 95% CI [0.00374, 0.0938]) compared to control participants (**Figure 3B1**). Additionally, shamanic practitioners had significantly greater connectivity in the low beta band during drumming (*t*(22.1) = 2.11, *p* = .047, 95% CI [0.000352, 0.0432]) and music (*t*(24.9) = 2.25, *p* = .033, 95% CI [0.00263, 0.0584]) compared to controls (**Figure 3B2**). These statistically significant measures of connectivity did not correlate with any of the OAV domains.

**Figure 3.**
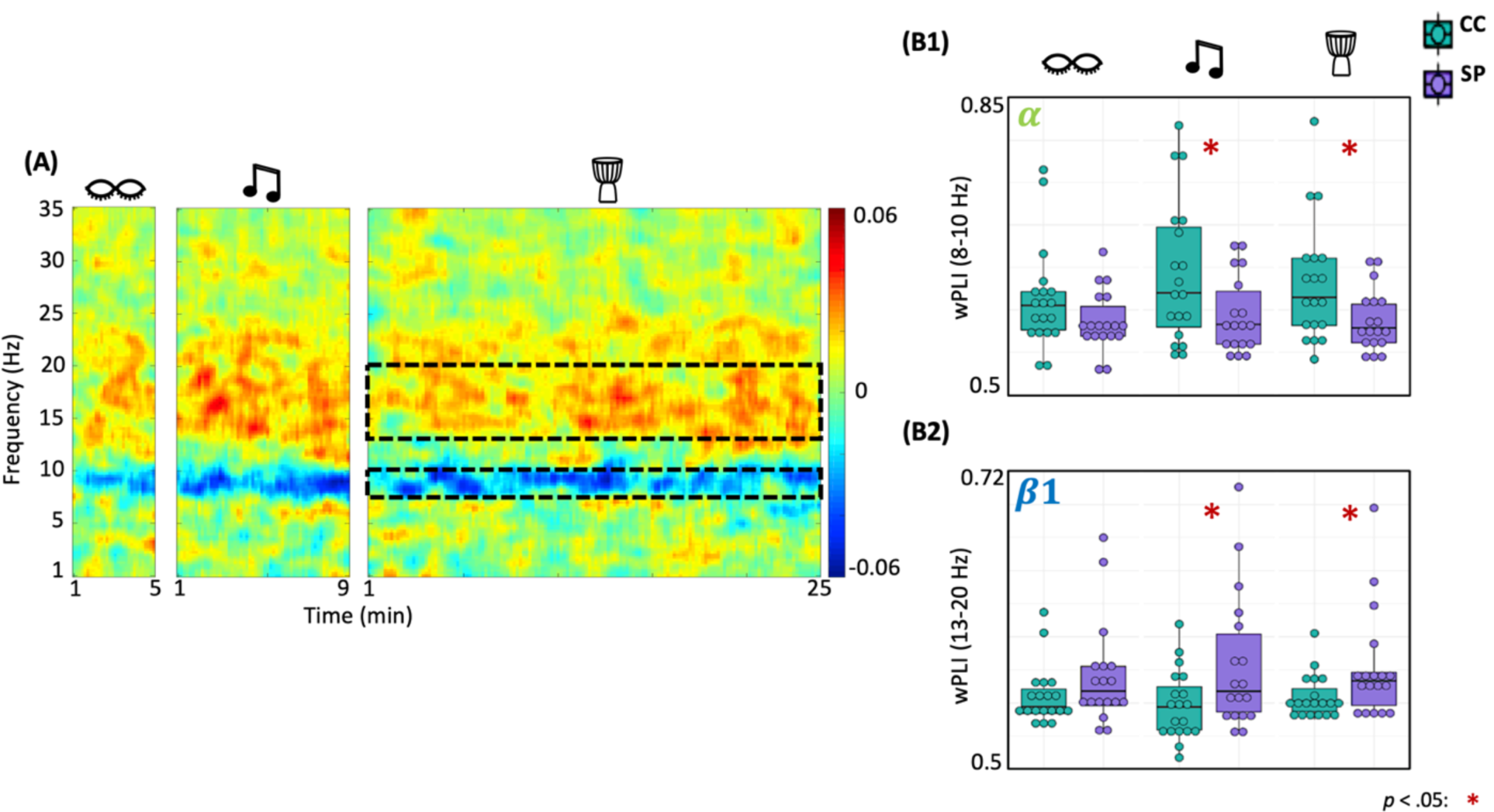
Shamanic practitioners have increased low beta and decreased low alpha connectivity during drumming. **(A)** Differences in functional connectivity, measured by weighted phase-lag index (wPLI), between shamanic practitioners (SP; n = 18) and control participants (CC; n = 19). Warmer colors represent greater connectivity in SP while cooler colors indicate greater connectivity in CC. The dotted lines represent the low alpha (8-10 Hz) and low beta (13-20 Hz) frequency bands, which were significantly different between groups. **(B1-2)** Box plots illustrating the average wPLI in the low alpha (**B1**) or low beta (**B2**) frequency bands in SP (purple) and CC (green) during eyes closed, drumming, or music. Whiskers represent the lower (25%) and upper (75%) quartiles and the center band of each box represents the median for the group. SP experienced statistically significant decreases in wPLI in the low alpha band during classical music and drumming compared to CC **(B1)**, as well statistically significant increases in the low beta band **(B2)**. Significance is indicated by the key in the lower righthand corner of the figure.

### Lempel-Ziv complexity in the gamma band decreases during drumming in shamanic practitioners and correlates with feelings of insightfulness

We evaluated differences in EEG signal diversity using Lempel-Ziv complexity (LZc) (**Figure 4A**). There was a significant main effect of group on LZc in the gamma band (*F*(1,35) = 4.79, *p* = .035). Post-hoc testing revealed that shamanic practitioners had decreased LZc in the gamma band during drumming (*t*(19.4) = -2.53, *p* = .02, 95% CI [-0.0274, -0.00262]) compared to control participants (**Figure 4B**). Further, LZc in the gamma band was negatively correlated with feelings of insightfulness (*r*_*s*_ = 0.5, *p* = .034)(**Figure 4C**). While analysis of broadband LZc revealed a statistically significant interaction between group and condition (Group x Condition: *F*(2, 70) = 3.7, *p* = .03), post-hoc t-tests did not reveal any significant differences between groups.

**Figure 4.**
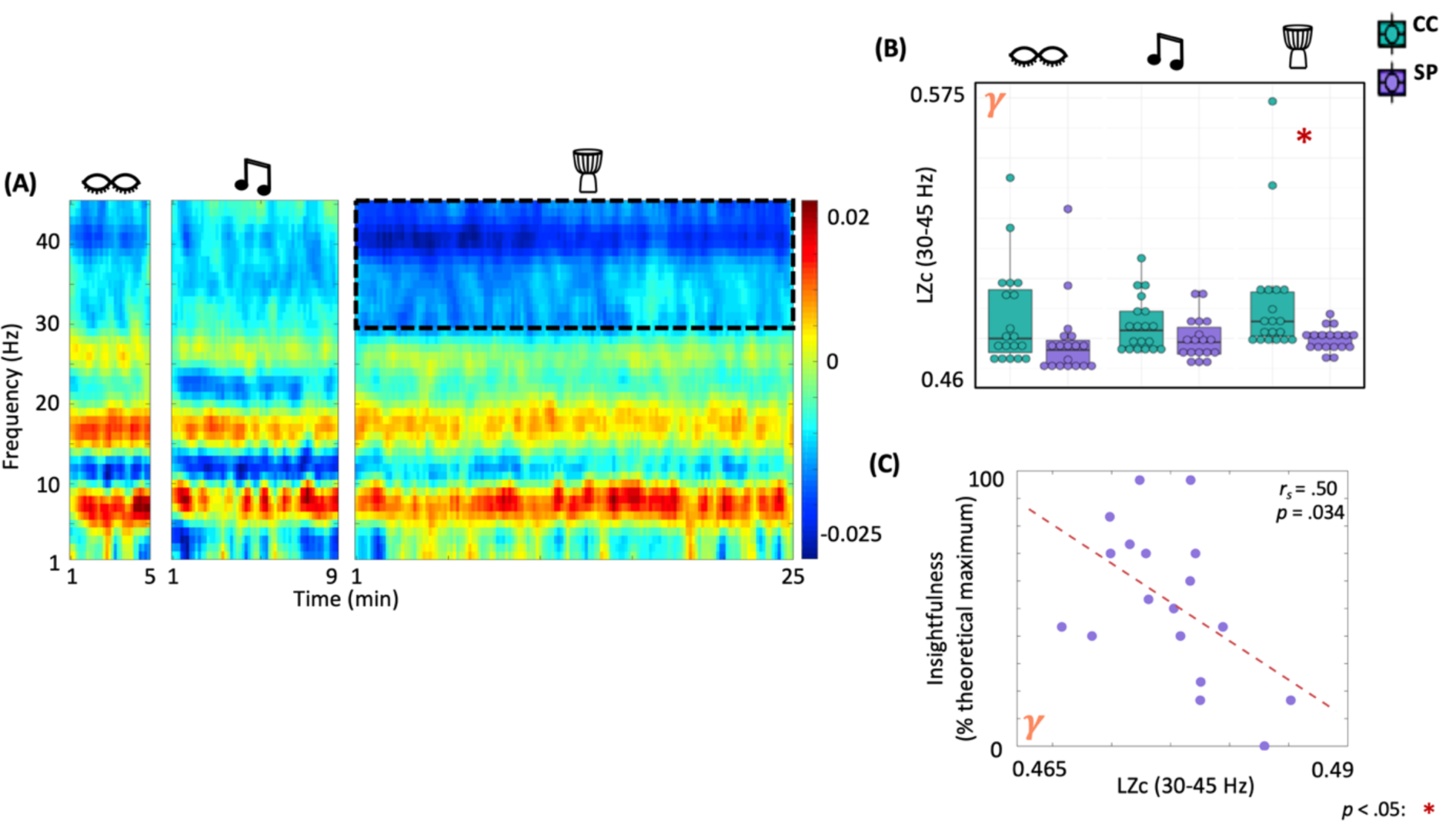
Shamanic practitioners have decreased neural signal diversity in the gamma band during drumming that is negatively correlated with feelings of insightfulness. **(A)** Comparison of the Lempel-Ziv complexity (LZc), a surrogate measure of neural signal diversity, across frequencies in shamanic practitioners (SP; n = 18) and controls (CC; n = 19). Warmer colors represent greater LZc in SP while cooler colors indicate greater LZc in CC. The dotted lines represent the gamma band (30-45Hz), which was significantly different between groups. **(B)** Box plots comparing the average LZc in the gamma band for SP (purple) and CC (green) during eyes closed, classical music, and drumming. Whiskers represent the lower (25%) and upper (75%) quartiles and the center band of each box represents the median for the group. SP experienced statistically significant decreases in LZc in the gamma band compared to CC. Significance is indicated by the key in the lower righthand corner of the figure. **(C)** Correlations between LZc in the gamma band and ASC domain scores revealed a negative correlation between gamma LZc and feelings of insightfulness.

### Shamanic practitioners experience increased criticality in beta and gamma bands during drumming that correlates with OAV domain scores

Finally, we examined differences in criticality using the pair correlation function (PCF)(**Figure 5A**) during eyes closed, music, and drumming. Given that the shamanic state of consciousness is characterized by a high level of mental activity (**Figure 1A**), we would anticipate increased PCF during shamanic trance, which would suggest increased criticality, and, in turn, increased metastability (i.e., patterns of global synchronization over time) and network susceptibility to internal and external perturbations. There was a main effect of group on PCF in the low beta (*F*(1,35) = 4.59, *p* = .039) and high beta bands (*F*(1,35) = 5.94, *p* = .02). For PCF in the gamma band, there were main effects of both group (*F*(1,35) = 10.09, *p* = .003) and condition (*F*(2,70) = 5.93, *p* = .004). Exploratory post-hoc t-tests revealed that, during drumming, shamanic practitioners had greater PCF in the low beta (*t*(34.5) = 3.32, *p* = .00213, 95% CI [0.00176, 0.00729])(**Figure 5B1**), high beta (*t*(33.2) = 3.29, *p* = .00235, 95% CI [0.00273, 0.0115])(**Figure 5B2**), and gamma (*t*(33.7) = 4.1, p < .001, 95% CI [0.00754, 0.0224])(**Figure 5B3**) bands compared to controls. Further, PCF in the low beta band positively correlated with the degree of complex imagery (*r*_*s*_ = .53, *p* = .023)(**Figure 5C1**) and elementary visual alterations (*r*_*s*_ = .56, *p* = .015)(**Figure 5C2**). Shamanic practitioners also had greater PCF in the high beta band during classical music (*t*(35) = 2.1, *p* = .043)(**Figure 5B2**) and the gamma band during classic music (*t*(31.5) = 2.75, *p* = .00987) and eyes closed (*t*(31) = 2.16, *p* = .039)(**Figure 5B3**).

**Figure 5.**
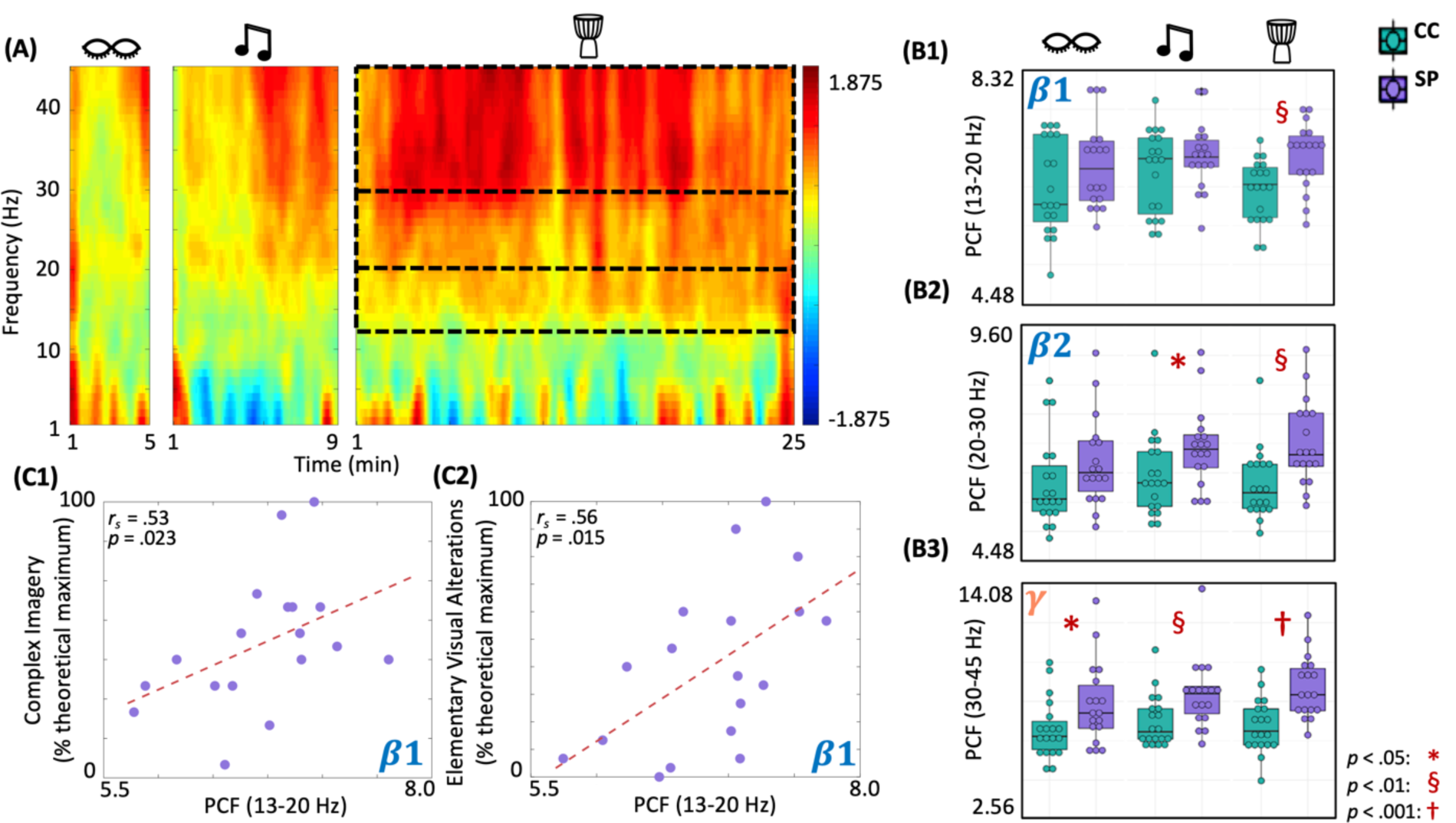
Shamanic practitioners have increased criticality in beta and gamma bands during drumming, with low beta increases correlating with complex imagery and elementary visual alterations. **(A)** Comparison of criticality, measured using a pair correlation function (PCF), across different frequencies between shamanic practitioners (SP; n = 18) and control participants (CC; n = 19). Warmer colors indicate greater PCF in SP while cooler colors represent increased PCF in CC. Dotted lines represent the low (13-20 Hz) and high (20-30 Hz) beta and gamma (30-45 Hz) bands, which were significantly different between SP and controls. **(B1-3)** Box plots illustrating the average PCF value for the low **(B1)** and high **(B2)** beta and gamma **(B3)** bands in SP (purple) and CC (green) during eyes closed, classical music, and drumming. Whiskers represent the lower (25%) and upper (75%) quartiles and the center band of each box represents the median for the group. SP had greater PCF in the low and high beta (**B1-2**) and gamma (**B3**) bands during drumming compared to CC, with increases extending to the classic music condition in high beta **(B2)** and gamma **(B3)** bands and the eyes closed condition in the gamma band **(B3**). Significance is indicated by the key in the lower righthand corner. **(C1-2)** Spearman correlations revealed a positive correlation between PCF in the low beta band and both complex imagery **(C1)** and elementary visual alterations **(C2)**.

## Discussion

### Summary

In the largest and most comprehensive neuroimaging study of shamanic practitioners to date, we characterized the shamanic state of consciousness using HD-EEG and a well-validated assessment for altered states of consciousness known as the OAV scale. We revealed shamanic practitioners to be significantly different from controls in a majority of the OAV domains during drumming. Further, these differences were similar in magnitude to individuals under the influence of ketamine, psilocybin, and MDMA. In addition to main effects of group on a variety of EEG measures, post-hoc exploratory analyses revealed that shamanic practitioners had increased absolute gamma power during drumming, which positively correlated with the degree of elementary visual alterations. Additionally, practitioners had decreased low alpha and increased low beta connectivity during drumming and classical music. Furthermore, signal diversity in the gamma band was decreased in shamanic practitioners during drumming, which negatively correlated with feelings of insightfulness. Finally, PCF was increased in the low beta, high beta, and gamma bands during drumming, with increases in the low beta band positively correlating with complex imagery and elementary visual alterations. Additionally, PCF in the high beta band was increased during classical music in shamanic practitioners compared to control participants, as was PCF in the gamma band during eyes closed and classical music. These findings suggest that, during shamanic trance, shamanic practitioners enter an altered state of consciousness that is accompanied by changes in a variety of EEG measures that correlate with changes in experience.

### The shamanic state of consciousness, gamma oscillations, and other absorptive states of consciousness

To our knowledge, the only previous investigation of shamanic trance using EEG was by Flor-Henry et al., who found increased absolute low (13-20 Hz) and high (21 – 50 Hz) beta power during shamanic drumming(9). These findings are similar to those of the current study, as we found increased gamma power (30 – 45 Hz) during drumming in shamanic practitioners compared to control participants. While we did not see differences in the low beta band, the presence of increased gamma power in both investigations supports the involvement of gamma oscillations in the shamanic state of consciousness. Gamma oscillations have been implicated in a variety of cognitive processes, including consciousness(20), maintenance of visual images within working memory(21), and visual perception(22), which is relevant given the correlation between gamma power and elementary visual alterations. Furthermore, previous investigations of brain activity during meditation have revealed increased gamma activity(23–28), with some attributing elevations in gamma to enhanced perceptual clarity(26), spontaneous visual imagery(29), or increased attentional effort(24). However, other trance states, such as possession trance, are frequently characterized by power changes in the theta, alpha, or beta bands(30–32), without analysis of the gamma band. Thus, while changes in the other frequency bands are absent during shamanic trance, it remains unknown whether our findings of increased gamma power are comparable. While it is difficult to pinpoint the exact reason of increased gamma power during shamanic trance, our findings are in agreement with previous literature on the shamanic state of consciousness and suggest similarities between shamanic trance and meditative states.

In addition to increases in gamma power, shamanic practitioners demonstrated decreased neural signal diversity and increased criticality (i.e., PCF) in the gamma band during drumming when compared to control participants. While studies of signal diversity and criticality in non-pharmacologically induced altered states of consciousness remain scarce, a recent magnetoencephalography (MEG) study during meditation assessed changes in entropy and metastability(33), which is directly related to criticality (e.g., increased metastability is associated with increased criticality). Martínez Vivot et al. revealed meditative practice to be characterized by increased entropy and decreased metastability in the gamma band(33). This is the inverse of our findings, where shamanic practitioners had *decreased* neural signal diversity and increased criticality in the gamma band, which, by definition, indicates *increased* metastability and brain network susceptibility. Given the paucity of research evaluating these measures in the context of shamanic trance or other absorptive states, it is difficult to discern the reason for this difference. One such explanation could be a difference in computational methods, as Martínez Vivot et al. evaluated changes in sample entropy (compared to Lempel-Ziv complexity) and utilized different techniques for measuring metastability. Additionally, there exist important differences in EEG and MEG, one being that MEG is more sensitive to tangential (but not radially-oriented) sources(34), which could potentially lead to discrepancies when comparing findings from these two methods. Furthermore, despite the similarities between shamanic journeying and deep meditative trance, they are distinct states of consciousness that are achieved through different means and for different purposes, meaning an inverse of findings may not be surprising. Future investigations characterizing non-pharmacologically induced altered states of consciousness and other absorptive or contemplative states are needed.

### Shamanic practitioners enter an altered state of consciousness similar yet distinct from the psychedelic state

Our analyses revealed significant differences between shamanic practitioners and control participants in 8 of the 11 OAV domains during drumming, with elevations in those 8 domains mirroring or exceeding domains altered during the psychedelic state. This suggests that shamanic practitioners are entering an altered state of consciousness during journeying and confirms the anecdotal overlap with the psychedelic state. However, the shamanic state of consciousness was different from the drug response in multiple OAV domains, suggesting that shamanic trance is a distinct altered state and does not perfectly correlate with the drug experience of any of the three substances examined by Studerus et al., even ones used for shamanic journeying (e.g., psilocybin). This assertion is supported by the lack of congruency between previous EEG and MEG findings during the psychedelic state and the results of the current study. Past studies have demonstrated widespread decreases in broadband power(35–40), with an emphasis on decreases in the alpha band that correlate with the degree of subjective effects(13, 39, 41, 42). Given our results demonstrating an overlap in reported changes in experience during the shamanic state of consciousness and the psychedelic state, it is surprising that shamanic practitioners not only lacked changes in alpha power, but experienced *increases* in power isolated to the gamma band that correlated with the degree of elementary visual alterations. Although increased gamma power has been reported during select studies of Ayahuasca exposure(43, 44)—an Amazonian brew commonly used for shamanic journeying that contains the psychoactive substance N, N-dimethyltryptamine—it was found to be unrelated to psychometric measures(44). Thus, it is possible that decreases in alpha power are specific to pharmacologic intervention of serotonergic psychedelics rather than changes in conscious experience, and that there are separate, distinct underlying neural mechanisms contributing to the subjective effects of psychedelics and the shamanic state of consciousness.

While literature investigating functional connectivity during the psychedelic state using EEG and MEG remains scarce, one study did find changes in lag phase synchronization in the delta band during psilocybin to be associated with feelings of insightfulness and spiritual experience(40). Our findings suggest that while shamanic practitioners differed from controls in low alpha and low beta connectivity, these changes are not associated with the degree of assessed altered states of consciousness.

Furthermore, changes in connectivity also occurred during classical music and, according to **Figure 3A**, to some degree at baseline, possibly reflecting long-term (i.e., trait) changes in low alpha and low beta connectivity in shamanic practitioners.

Previous literature examining changes in neural signal diversity during the psychedelic state have consistently revealed increases in Lempel-Ziv complexity(45, 46) that correlate with the strength of psychedelic-induced changes in phenomenal content(45, 46), particularly in visual domains. Yet, we found that the shamanic state of consciousness was characterized by *decreases* in neural signal diversity in the gamma band that negatively correlated with feelings of insightfulness, suggesting that decreased diversity in the neural signal was indicative of a more profound experience. One possible explanation for this discrepancy is that previous literature has focused on broadband signal diversity rather diversity within a given frequency band. However, analysis of broadband LZc did not reveal any significant differences between groups. When coupled with the lack of decreased alpha power, our findings suggest that while shamanic trance may be anecdotally similar to the psychedelic state, they are discrete altered states of consciousness, most likely with different underlying mechanisms by which changes in the subjective experience originate. This is further supported by previous literature demonstrating increased activity in the default-mode network (DMN) during shamanic journeying(8) rather than the characteristic decreases found during the psychedelic state(39, 47, 48), with decreases in alpha power that correlated with feelings of ego dissolution localizing to nodes of the DMN(38). Future studies should aim to investigate shamanic trance with and without the use of pharmacological intervention to further characterize the relationship between shamanic and psychedelic states.

### The shamanic state of consciousness and the entropic brain hypothesis

According to the entropic brain hypothesis, consciousness arises within a critical zone rather than a critical point(49), which is supported by a series of studies demonstrating that a hierarchical modular structure of the brain network extends a critical point to a critical region(50). The extended critical region allows for increased sustainability, which in turn enables the brain to maintain criticality under perturbations. Furthermore, the entropic brain hypothesis suggests that normal waking consciousness exists within a band positioned in the middle of this critical zone and changes in consciousness occur when pushed beyond the lower or upper limits of this band. For example, sedatives and anesthetics shift the brain downward, closer to a lower limit (i.e., sub-critical or extremely ordered) diminishing conscious contents. On the contrary, psychedelics shift the brain upward, closer to the upper limit of the critical zone (i.e., super-critical or extremely disordered) with richer conscious contents(49). This was recently demonstrated by Atasoy et al., who provided empirical evidence for heightened criticality during Lysergic acid diethylamide (LSD) using human fMRI blood oxygen level dependent (BOLD) signals(51).

In the current study, we found that shamanic trance increased PCF in beta and gamma bands. As mentioned previously, increased PCF is indicative of increased criticality, and in turn increased brain network susceptibility to perturbations and metastability. Based on the assumption that the brain during normal waking consciousness resides near criticality, increased PCF of shamanic practitioners compared to the control group implies the shamanic state of consciousness shifts the brain state to the upper limit of the critical zone. With heightened criticality, the brains of shamanic practitioners may be more susceptible to internal and external perturbations, which produce more diverse spatiotemporal brain activity patterns and consequently allow for a “richer” or more “diverse” conscious experience. This is supported by the correlations between increased PCF in the low beta band and the ASC domains of complex imagery and elementary visual alterations. Furthermore, while PCF increases were greatest during shamanic drumming, PCF was also increased during classical music and eye closure in shamanic practitioners compared to the control group, suggesting that shamanic journeying may induce enduring changes in criticality, shifting the brain to reside in the upper bounds of the critical zone long-term (i.e., over the course of months or years). Future longitudinal studies could expand these findings by examining criticality and other EEG measures at various points within the lifetime of shamanic practitioners.

### Methodologic Strengths and Weaknesses

This study has many strengths, as well as several notable limitations. In addition to validating the spectral changes noted in the only previous EEG study of shamanic trance, we characterized the shamanic state of consciousness using a variety of computational methods used previously to assess other altered states of consciousness. Furthermore, we investigated the relationship between such measures and changes in subjective experience, as reported by shamanic practitioners. This allowed for the most comprehensive characterization of the shamanic state of consciousness to date, using the largest cohort of shamanic practitioners reported in the literature. However, our sample size is small relative to many neuroimaging studies. This led to the lack of adjustment for multiple comparisons in our post-hoc analyses due to their exploratory nature. Yet, it is important to note that trend relationships (*p* = .052 - .07) would remain for a majority of EEG measures (connectivity, signal diversity) and almost all PCF results, excluding in the high beta band during music (*p* = .065), would remain significant (*p* < .05) after correction (Benjamini-Hochberg procedure).

In order to allow for greater generalizability of findings and ease of recruitment, our cohort of shamanic practitioners had diverse backgrounds of shamanic practice and rituals. While considered a strength of this study, this also introduced potential variability in our study population. Additionally, our control participants were asked to rest quietly during the shamanic drumming period while our shamanic practitioners were asked to enter a shamanic trance. This means shamanic practitioners were cognitively engaged during this time while controls were passively listening, which could have led to a difference in EEG changes. However, unlike previous studies of the shamanic state of consciousness, we collected data during classical music as a contrast condition in order to identify brain activity specific to shamanic trance rather than due to cognitive engagement.

We did not collect electromyography data to control for movement, meaning it remains possible that our findings within the gamma band are contaminated by muscle artifact. However, we did not anticipate any differences in muscle contamination across conditions or experimental groups, meaning any differences within the gamma band can likely be attributed to changes in brain activity rather than movement.

### Conclusion

We found that shamanic practitioners entered an altered state of consciousness compared to control participants during shamanic drumming, with the magnitude of these subjective experiences being comparable to reports of the psychedelic state. Furthermore, shamanic practitioners evinced changes in a variety of EEG measures during shamanic drumming that correlated with measures of their altered states of consciousness. These findings suggest that shamanic trance and psychedelic drug-induced altered states of consciousness are “distinct states with shared traits,” meaning that despite various overlapping phenomenal components, they are characterized by different changes in brain activity. This suggests the involvement of endogenous mechanisms for non-pharmacologically-induced altered states of consciousness that are distinct from those co-opted by psychedelic drugs.

## Materials and Methods

### Ethics

This study was approved by the University of Michigan Institutional Review Board (HUM00089064). Written informed consent was obtained from all participants prior to study procedures.

### Participants

Two separate populations were recruited for this study: 24 experienced shamanic healing practitioners and 24 age- and sex-matched control participants. Shamanic practitioner inclusion criteria was as follows: five or more years of practice in shamanic techniques; performed a minimum of 40 shamanic healing sessions over the last 5 years, primarily focused on healing individual people; received training in at least one shamanic tradition or style under expert supervision; used rhythmic drumming during shamanic journeying and healing practice; and possessed the ability to enter a shamanic trance in 15 minutes and complete shamanic healing session in 30 minutes while sitting still and quiet. Practitioners were excluded if they had a history of traumatic brain injury or epilepsy. Control participants were age- and sex-matched to shamanic practitioners with the following inclusion criteria: no current or past experience or interest in shamanic practice or media involving shamanism; no current or past history of meditation, spiritual healing, or spiritual training; no current or past history of hallucinogen, entheogen, or psychedelic drug use; no current or past experience with trance or hypnosis, or interest relating to media of such themes; no current or past history of being a drummer, frequent listening to drumming music, or a history of professional musicianship. Control participants were excluded if they had current or past diagnosis of depression, anxiety, schizophrenia, or other serious psychological disorders, untreated major physical or psychological illness, a history of traumatic brain injury, or a history of epilepsy. Subjects were excluded from analyses for technical difficulties pertaining to EEG acquisition (n = 4), EEG data quality issues (n = 4), or lack of questionnaire data (n = 2), resulting in 18 shamanic practitioners and 19 control participants in the final cohort for analysis. Demographics for these participants are illustrated in **Supplemental Table 2**.

### Pre-Study Procedures

Prior to their study visit, shamanic practitioners were asked to practice journeying during the 25 minutes of rhythmic drumming music provided during the study visit(52), while control participants were asked to listen to the same pre-recorded drumming without any attempts at journeying.

### Study Design

Participants underwent baseline EEG recordings with their eyes open for five minutes, followed by five minutes of eye closure. Following baseline recordings, each participant underwent three experimental blocks: cognitive testing, shamanic drumming, and classical music listening. The order of blocks was counterbalanced between groups. Participants were asked to rest again with their eyes opened and closed for a 5-minute period following the last experiment. EEG data was collected continuously throughout all experiments. Example study day timelines are illustrated in **Figure 6A**. Given that this manuscript is centered on brain activity during the shamanic state of consciousness, cognitive testing results will not be discussed.

**Figure 6.**
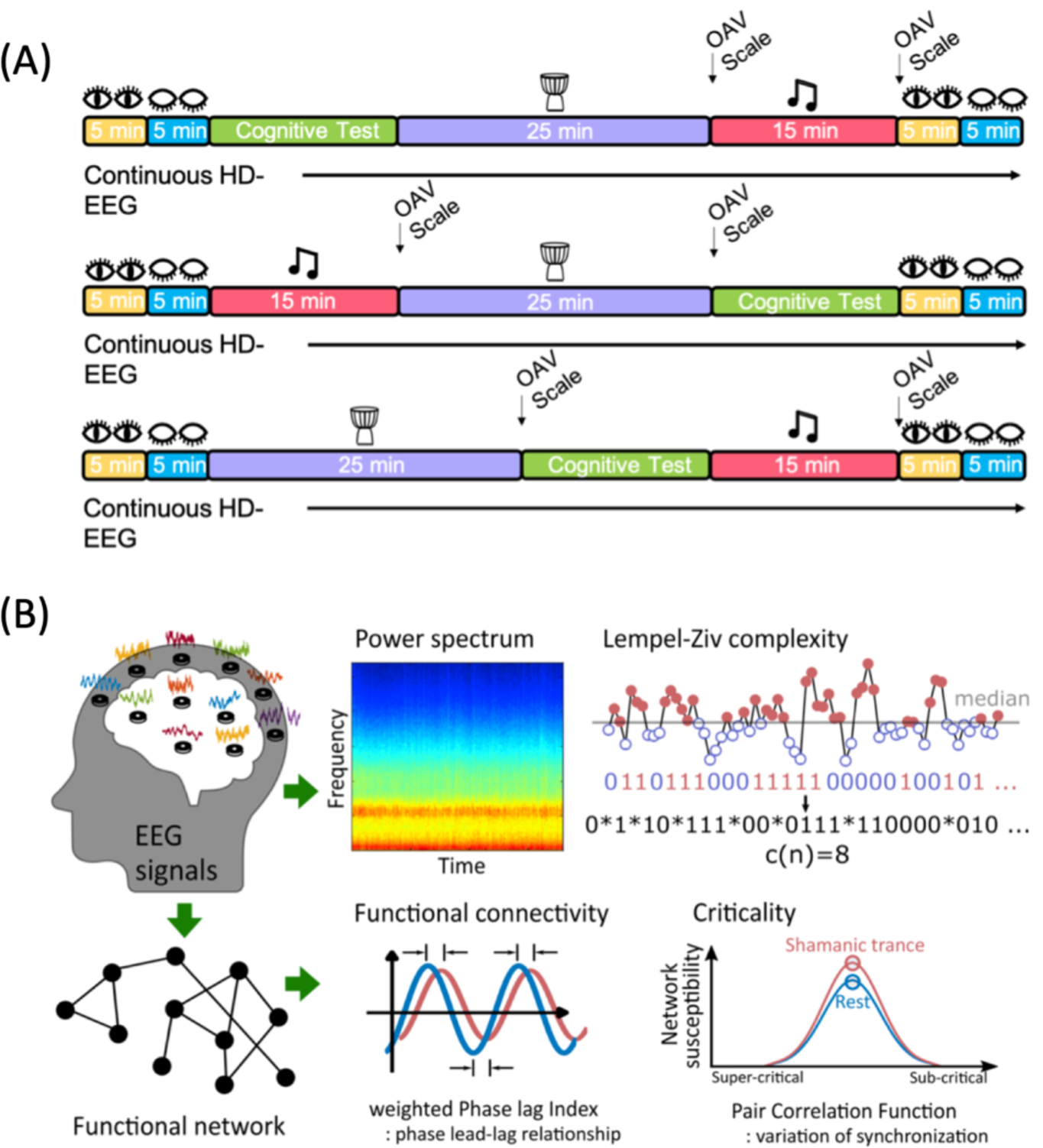
Study design. **(A)** Example timelines of a typical study day based on a given experiment block. Each individual was randomized to the order they underwent cognitive testing, shamanic drumming, and classical music listening. Eyes open and eyes closed resting periods occurred at the beginning and end of the experimental day. The Altered States of Consciousness Questionnaire, known as the OAV scale, was administered following classical music and shamanic drumming. **(B)** All participants underwent high-density electroencephalogram (HD-EEG) recordings during drumming, classical music, and periods of eye opening and closure. Following preprocessing, HD-EEG signals were used to compute subsequent measures within each frequency band, including the power spectrum, functional connectivity measured by weighted Phase Lag Index, signal diversity measured by Lempel-Ziv complexity (image referenced from Leemburg & Bassetti, 2018), and criticality measured by the pair correlation function.

### Shamanic Healing Experiment

During the shamanic healing experiment, shamanic practitioners and controls listened to the same pre-recorded drumming they listened to at home prior to the study visit. This recording consisted of rhythmic drumming accompanied by a mixture of bells and rattles and combined a 15-minute drumming piece with the middle 10 minutes of this piece repeated, for a total of 25 minutes(52). Shamanic practitioners were asked to conduct shamanic healing during this time, during which they entered an altered state of consciousness in order to glean information to be used for the physical, psychological, or spiritual healing of a client not present during the study. Control participants were instructed to rest quietly and both groups were asked to keep their eyes closed during the drumming period. Following the drumming period, shamanic practitioners and controls completed an Altered States of Consciousness (ASC) questionnaire known as the OAV scale(53), which contains 66 items pertaining to their subjective experience.

### Classical Music Experiment

The classical music experiment consisted of 15 minutes of classic music listening, which consisted of two songs (piano and violin) played in succession(54)(55). Both shamanic practitioners and control participants were asked to listen with their eyes closed during this time. Participants again filled out the OAV scale following classical music listening.

### Qualitative Analysis

The 66-item OAV Scale questionnaire data were pooled into 11 domains according to previous literature(19), including experience of unity, spiritual experience, blissful state, insightfulness, disembodiment, impaired control and cognition, anxiety, complex imagery, elementary visual alterations, audio-visual synesthesia, and changed meaning of percepts. The percent of the theoretical scale maximum was calculated for each domain for each individual, as well as the mean and standard error of the mean (SEM).

### EEG Acquisition

Participants were fitted with a 129-channel HydroCel Geodesic Sensor Net (Electrical Geodesics, Inc./Phillips Eugene, OR, USA) according to head circumference. EEG data were acquired at a 500-1000 Hz sampling rate with a vertex reference using NetAmps 400 amplifier and Netstation 4.5 software. Channel impedances were reduced to below 50 kΩ per the manufacturer’s instruction.

### EEG Analysis

Using the MATLAB open source toolbox EEGLab(56), data were visually inspected for artifact and preprocessed via resampling to 500 Hz as necessary, bandpass filtering between 1 – 45 Hz, and re-referencing to the global average. Data were epoched into 10-second non-overlapping windows; all analyses were conducted on sequential 10-second windows. Subsequent analyses focused on absolute power, functional connectivity, neural signal diversity, and criticality (**Figure 6B**) within each of the following frequency bands: delta 1 – 4 Hz, theta 4 – 8 Hz, low alpha 8 –10 Hz, high alpha 10 – 13 Hz, low beta 13 – 20 Hz, high beta 20 – 30 Hz, and gamma 30 – 45 Hz (except for connectivity, which was limited to 30 – 35 Hz to ensure reliability of this measure). While we did not collect electromyography data to control for muscle artifact, we did not anticipate differences in muscle contamination across participants and conditions, meaning any differences in the gamma frequency band were presumably from electroencephalographic differences. EEG measures were computed for the full duration of periods of eye closure (5 minutes) and shamanic drumming (25 minutes). Analyses of brain activity during classical music were performed on the first song period (∼10 minutes).

### Spectral Analysis

Spectral power was calculated using the ‘spectrogram.m’ function in the MATLAB Signal Processing Toolbox (time window: 3s hamming window, overlap: 50%) based on the short-time Fourier transform. For each state (eyes closed, classical music, drumming), absolute power within each frequency band was averaged across all time windows and channels.

### Functional Connectivity

Functional connectivity was assessed using the average weighted Phase Lag Index (wPLI), which measures the phase lead-lag relationship between two signals. If two signals are found to be functionally coupled within a given frequency band, the resulting wPLI value will be high within that band. wPLI is robust to the EEG volume conduction problem(57, 58) and improves Phase Lag Index (PLI) with weights of the imaginary part of the cross-spectrum, reducing noise and volume conduction effects. When the imaginary part of the cross-spectrum is ℑ[X],

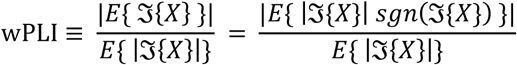

where the signed PLI is the numerator normalized by denominator, the imaginary part of the cross-spectrum. The wPLI was calculated for each frequency band and averaged across time and all channels in each state.

### Neural Signal Diversity

Recent literature examining psychedelic-induced altered states of consciousness have sought to characterize changes in signal diversity(13, 46), which reflects the number of unique (i.e., diverse) patterns within a signal. Given the aforementioned overlap between the psychedelic state and the shamanic state of consciousness, we measured and compared neural signal diversity of each state using Lempel-Ziv complexity (LZc), which calculates the temporal complexity of the signal by computing the number of distinct subsequences through the whole signal(59) (**Figure 6B**, image referenced from Leemburg & Bassetti, 2018(60)). The time series of each EEG epoch was binarized using the median value (M) of each time series(60, 61) as a threshold.

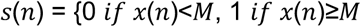

where *x*(*n*) is each data point of the original time series and the median value is M. The median value of each epoch is used for the robustness to outliers and the invariance from the signal amplitude.

A new subsequence of consecutive numbers (“words”) was counted as *c*(*n*) through the binarized time series and normalized by the theoretical maximum number of words b(*n*)=*n*log_2_*n*.

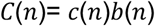

LZc was computed for each frequency band in each state and averaged across time and channels. To better compare changes in LZc during shamanic trance to those in previous literature (i.e., psychedelic states), we computed LZc of the broadband (1-45Hz) signal, as well as within individual frequency bands.

### Criticality

Criticality, a balanced state between order and disorder(62), is observed ubiquitously in physical and biological systems and is accompanied by various functional benefits, including large information storage, optimal information transmission, and flexible response to external stimuli(63). Many computational and experimental studies suggest that human and animal brains reside near a critical point during conscious wakefulness(64) and a loss of criticality induces altered states of consciousness such as sleep, anesthesia, and unresponsive wakefulness syndrome(15–18). We assessed criticality using the pair correlation function (PCF), which is the variance of network synchronization, reflecting network susceptibility to internal and external perturbations (e.g., from stimuli)(65). PCF is maximal at a critical point and gradually decreases as a system moves toward a sub- or super-critical state(66). Recently, the reliability of PCF in evaluation of the level of consciousness was supported with a computational model and EEG during general anesthesia(66).

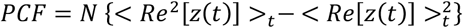

where Re[*z(t)*] is the real part of the *z(t)*.

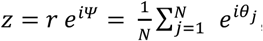

where Ψ is the order parameter phase. The absolute value *r* = |z| represents the degree of synchronization. The *r* is equal to zero when the phases of nodes are uniformly distributed and one when all the nodes have the same phase.

### Statistical Analysis

For comparison of OAV domain scores, Wilcoxon signed-rank tests were used to assess differences in paired samples and Wilcoxon rank-sum tests were used to assess differences between shamanic practitioners and controls. With permission from Studerus et al.(19) and using open-access data, we used the mean and standard deviation to compare OAV domain scores of shamanic practitioners and individuals under the influence of psychedelic substances using unpaired t-tests. A two-way repeated measures analysis of variance (ANOVA) was used to assess differences in EEG measures between groups (shamanic practitioner vs. control; between-subject factor) and across conditions (eyes closed vs. classical music vs. drumming; within-subject factor), as well as any interaction between these two factors (group x condition). Exploratory post-hoc analysis of group differences within each condition were conducted using unpaired t-tests. Given our small sample size and the exploratory nature of post-hoc tests, we did not adjust for multiple comparisons. Spearman correlations were conducted to assess relationships between EEG measures during drumming that were statistically significantly different between groups and OAV domain scores in shamanic practitioners. Findings were considered statistically significant if *p* < .05.

## Data availability

All data and code used for this project will be made available upon request of the authors with a data sharing agreement between institutions.

## Acknowledgements

We would like to thank Dr. Erich Studerus for his agreement to and helpful suggestions for comparing our data from shamanic practitioners to healthy volunteers under the influence of psychedelics. Additionally, we would like to thank Kate Durda and Stephanie Tighe, who offered invaluable insight as shamanic practitioners when designing our study. This work was funded by the Center for Consciousness Science, Department of Anesthesiology, University of Michigan Medical School.

