## Supplemental Table 1 for "Neural correlates of the shamanic state of consciousness"

**Supplemental Table 1. Statistical comparison of OAV domain scores between shamanic practitioners and healthy individuals under the influence of psychedelics (Studerus et al., 2010).**

| | Ketamine (df = 178) | | | Psilocybin (df = 343) | | | MDMA (df = 118) | | |
|---|---|---|---|---|---|---|---|---|---|
| **OAV Domain** | *t* | *p* | 95% CI | *t* | *p* | 95% CI | *t* | *p* | 95% CI |
| Complex imagery | **4** | **<.001** | **[14.57, 42.88]** | **2.68** | **.0077** | **[5.54, 36.09]** | **7.95** | **<.001** | **[34.86, 58.002]** |
| Exp. of unity | **2.83** | **.0052** | **[6.27, 35.12]** | **3.8** | **<.001** | **[13.05, 41.06]** | **4.11** | **<.001** | **[15.64, 44.66]** |
| Spiritual exp. | **6.43** | **<.001** | **[26.17, 49.33]** | **6.72** | **<.001** | **[27.65, 50.55]** | **9.34** | **<.001** | **[36.59, 56.27]** |
| Blissful state | **4.51** | **<.001** | **[16.75, 42.82]** | **2.93** | **.0036** | **[6.66, 33.91]** | 1.5 | .14 | [-3.6, 25.98] |
| Disembodiment | .737 | .46 | [-9.7, 21.26] | **3.96** | **<.001** | **[13.74, 40.90]** | **4.37** | **<.001** | **[16.28, 43.32]** |
| Insightfulness | **4.83** | **<.001** | **[17.3, 41.21]** | **4.44** | **<.001** | **[15.23, 39.41]** | **6.19** | **<.001** | **[25.03, 48.57]** |
| Elem. visual alt. | 1.1 | .28 | [-6.64, 23.21] | 1.11 | .27 | [-24.77, 6.91] | **7.35** | **<.001** | **[25.28, 43.93]** |
| Changed percepts | .099 | .92 | [-12.88, 14.25] | .64 | .52 | [-19.62, 9.99] | 0.48 | .63 | [-11.52, 18.94] |
| Audio-visual syn. | **2.17** | **.031** | **[-32.94, -1.57]** | **3.16** | **.0017** | **[-42.45, -9.85]** | 0.045 | .96 | [-10.22, 9.77] |
| Imp. cont. & cogn. | **3.88** | **<.001** | **[-35.8, -11.65]** | **2.86** | **.0045** | **[-24.62, -4.55]** | 1.41 | .16 | [-16.91, 2.86] |
| Anxiety | **2.1** | **.037** | **[-17.82, -0.56]** | 1.12 | .27 | [-12.96, 3.57] | .012 | .99 | [-5.39, 5.33] |

df = degrees of freedom; *t* = test statistic; *p* = significance; CI = confidence interval [lower, upper]
