## Supplemental Table 2 for "Neural correlates of the shamanic state of consciousness"

**Supplemental Table 2. Demographics of shamanic practitioners and controls included in analysis.**

| | Shamanic Practitioners (n = 18 | Controls (n = 19) |
|---|---|---|
| **Age (median)** | 56 (5.5) | 57(12) |
| **Sex (% male)** | 50% | 47% |
| **Years of Practice** | 19.5(21.125) | --- |
| **Healing Sessions Per Month** | 12.75(14.25) | --- |
